# The adhesion capability of *Staphylococcus aureus* cells is heterogeneously distributed over the cell envelope

**DOI:** 10.1101/2021.01.05.425282

**Authors:** Christian Spengler, Erik Maikranz, Bernhard Glatz, Michael Andreas Klatt, Hannah Heintz, Markus Bischoff, Ludger Santen, Andreas Fery, Karin Jacobs

## Abstract

Understanding and controlling microbial adhesion is an important biomedical problem. However, many properties of the adhesion process of bacteria are still unknown, for example the distribution of adhesive strength over the cell wall. While a patchy colloid model for adhesion has been developed recently for Gram-negative *Escherichia coli* cells, a comparable model for Grampositive cells is unknown. Here, we use single-cell force spectroscopy to measure the adhesion of *Staphylococcus aureus* at different positions on tailored surfaces. We find heterogeneous adhesion profiles with varying degrees of intensity. By comparing these results to simulations, we find that locally increased adhesion can be explained by several distinct spots of high adhesion capabilities, similar to the patchy colloid model. Only for the underlying profile without local adhesive spots simple geometric considerations are insufficient. Rather, strong angle-dependent molecule-substratum interactions are necessary to explain the bathtub-like adhesion profiles seen for *Staphylococcus aureus* on a sinusoidal surface. We discuss implications of our results for the development of new materials and the design and analysis of future studies.

---

Infections caused by bacterial biofilms are a major healthcare problem^1–3^. These biofilms can be found both on natural surfaces, e. g. in the nasal^4^ and oral^5^ cavity, as well as on artificial surfaces, such as the exterior of prostheses, catheters and other medical devices^6–9^. In this context, *Staphylococcus au-reus* (*S. aureus*) is an important human pathogen^10,11^, which is capable of forming biofilms with increased resistance to antibiotic treatment^12^ and the body’s own immune system^13^. Consequently, *S. aureus* can cause various diseases^14^, such as superficial skin disease, sepsis, endocarditis and pneumonia and numerous implant-associated infections^10^. Since the formation of a biofilm begins with the attachment of single bacterial cells, understanding and controlling bacterial adhesion to solid surfaces is an urgent challenge in biomedical research.

Previous studies demonstrated that *S. aureus* cells adhere by tethering cell wall macromolecules, the number of which varies greatly depending on the properties of the underlying substrate^15,16^. The number and properties of individual tethering molecules define the adhesive strength^17^, and by length fluctuations, the molecules can overcome certain degrees of surface roughness^18^. For the secretion and deposition of adhesins on the *S. aureus* cell wall, different mechanisms have been unraveled^19^. In particular, it has been shown that protein A is secreted very selectively near the septum and then built into the cell wall^20^. In another study, however, accumulation of protein A was also observed in additional areas of the cell wall and differences in the frequency and density of these clusters depending on the growth phase could be detected^21^. The same study also observed cluster formation for clumping factor A (ClfA), whose size, but not the frequency, was growth phase-dependent^21^. Atomic force microscopy (AFM) has been used in many studies to find specific interactions between functionalized probes and certain proteins at the cell wall^22–27^. While in these studies, ClfA and B as well as the fibronectin-binding protein A (FnbpA) have not been found to be distributed in distinct clusters^23–25^, it has been found that the collagen-binding protein (Cna) in *S. aureus*^26^ and Serine-aspartate repeat-containing protein G (SdrG) in *Staphylococcus epidermis*^27^ show a cluster-like distribution. Furthermore, a recent study utilizing DNA-PAINT, showed that in *S. aureus* the density of fibronectin-binding proteins is so small, that their interaction with flat surfaces is limited to the binding of single heterogeneously distributed molecules^28§^. Recently, mechanisms that can lead to protein clustering in lipid membranes have been deciphered by single-molecule atomic force microscopy^29^. However, the question of how the overall adhesion capability of *S. aureus* or other Gram-positive cells is distributed over the cell surface has not yet been answered. For Gram-negative *Escherichia coli* (*E. coli*) cells, it has been found recently that this species adheres to glass surfaces by adhesive patches on their cell wall, and that the number of patches defines the adhesive strength^30^. However, Gram-positive *S. aureus* cells have a very different cell wall composition and cell division behavior than *E. coli* cells, and it has been shown that the size of the contact area between cell and surface does not correlate with its adhesive strength. In particular it has been shown that the size of the contact area of different strains is largely comparable while they may differ vastly in terms of adhesive strengths^17^.

In this study, we performed single-cell force spectroscopy (SCFS) on a periodically structured surface to directly measure the adhesion of different positions of the *S. aureus* cell wall to an unconditioned abiotic surface. We used a polydimethylsiloxane (PDMS) substrate with a symmetrical, periodic surface topography with a wavelength in the size range slightly above the cell diameter. The surfaces were formed by a controlled wrinkling process that allowed patterning in a scalable fashion and has found applications in various studies^31–33^. With these substrates and the precise control of the lateral distance between several consecutive force-distance curves on the substrate’s surface, we were able to probe different parts of the cell envelope in terms of their adhesion capability. Our experiments show that the adhesive strength at a given position is quite robust over the course of several measurements but can vary greatly for different cell wall positions depending on the individual cell. By reproducing these experimental results with Monte Carlo (MC) simulations by extending a recent model of adhesion^15^, we show the importance of angle-dependent molecule-substratum interactions and that the adhesion capability of *S. aureus* is driven by adhesins organized in distinct patches of various number over the cell envelope. These results are important for the fabrication of new materials and the design of more precise models to describe bacterial adhesion.

## 1 Results and Discussion

### 1.1 Periodically Wrinkled PDMS Surfaces as Suitable Substrates

Since AFM-based force-distance curves can only be recorded by a vertical movement of a bacterial probe, surfaces providing flanks with slopes of suitable absolute values in positive and negative direction are required to measure the adhesion at different positions of the cell surface by SCFS (see Fig. 1a). Moreover, a substrate with a continuous transition from positive to negative local slopes would allow to probe not only two points, but also intermediate positions (see Fig. 1b). These requirements can be met by wrinkled PDMS surfaces, which are shown in Figure 1c and 1d^31,32^.

**Fig. 1.**
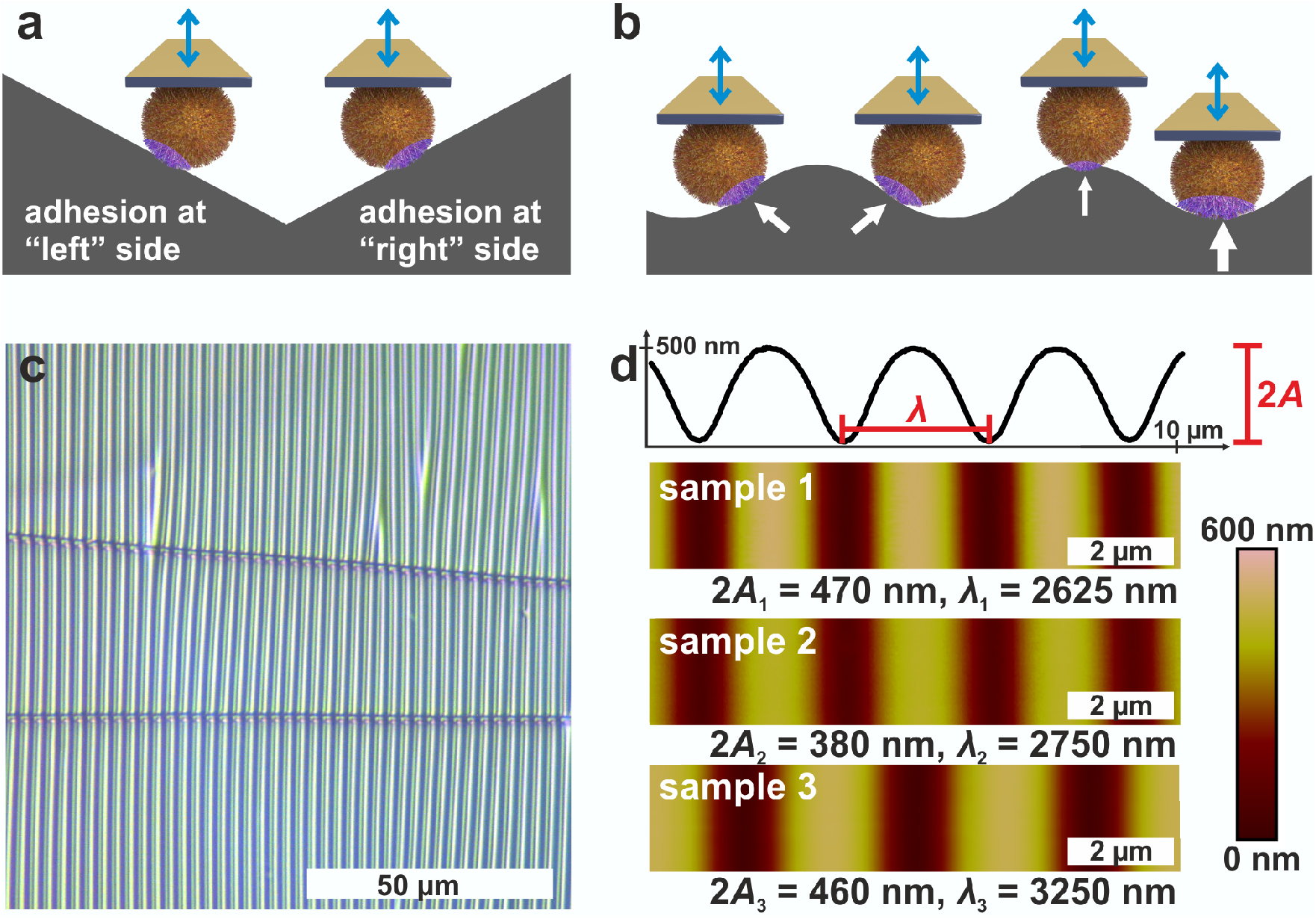
Measuring principle on wrinkled PDMS. a) Probing both sides of a cell on a substrate with negative and positive slope. b) Possibility to continuously probe different positions (indicated by arrows) of a cell on a substrate with a symmetric structure of changing local slope. c) Optical micrograph of wrinkled PDMS. d) AFM images of the wrinkled PDMS surfaces. For sample 1, a cross-section of the surface is shown to define the structure’s amplitude *A* and wavelength *λ*, which are displayed for every sample surface.

The optical micrograph shows that the wrinkled PDMS substrate has quite a homogeneous wrinkle structure over a large area, which is only rarely disrupted by cracks in the material (see Fig. 1c). To characterize the surface topography in all dimensions, the wrinkled PDMS was analyzed via topographical AFM (see Fig. 1d). For our experiments, we used three different PDMS samples, which were produced with slightly varying parameters. Figure 1d shows AFM images of all samples. In addition, a scan line recorded on sample 1 is depicted, in which the specific wrinkling parameters, wavelength *λ* and amplitude *A*, are defined. All samples have a very homogeneous surface structure: Locally and parallel to the trenches, the surface is very smooth with a root mean square roughness calculated parallel to the trenches (i.e. in y-direction in Fig. 1) below 5 nm. Perpendicular to the trenches, all surfaces feature a nearly sinusoidal periodicity that results in a vigorously homogeneous surface and a high symmetry within its repetitive structuring. The wavelengths and amplitudes of the periodic structures are in the same size range as the dimensions of *S. aureus* cells, as sketched in Figure 1b. Therefore, the wrinkled PDMS surfaces are a well-suited substrate to determine the adhesion force of *S. aureus* cells at different locations on the cell wall by SCFS.

### 1.2 Periodic Adhesion Patterns of *S. aureus* on Wrinkled PDMS - Construction of Adhesion Profiles

To measure the adhesion of *S. aureus* cells at different positions relative to the periods of the wrinkled PDMS surfaces, the substrates were mounted in a way that the trenches on the surfaces were parallel to the y-direction of the AFM scan area. Correct positioning was verified by scanning the surface before performing SCFS experiments. (An inclination of up to 1^*°*^ was accepted, otherwise the sample was repositioned.)

Then, several hundred force-distance curves were recorded with one and the same single cell while the x-position between each two consecutive curves was changed by a constant value (of 20-30 nm). From every curve, the adhesion force and the z-height at which the retraction began (termed “initial retraction height”) were determined, and the results are shown in Figure 2a for one exemplary cell.

**Fig. 2.**
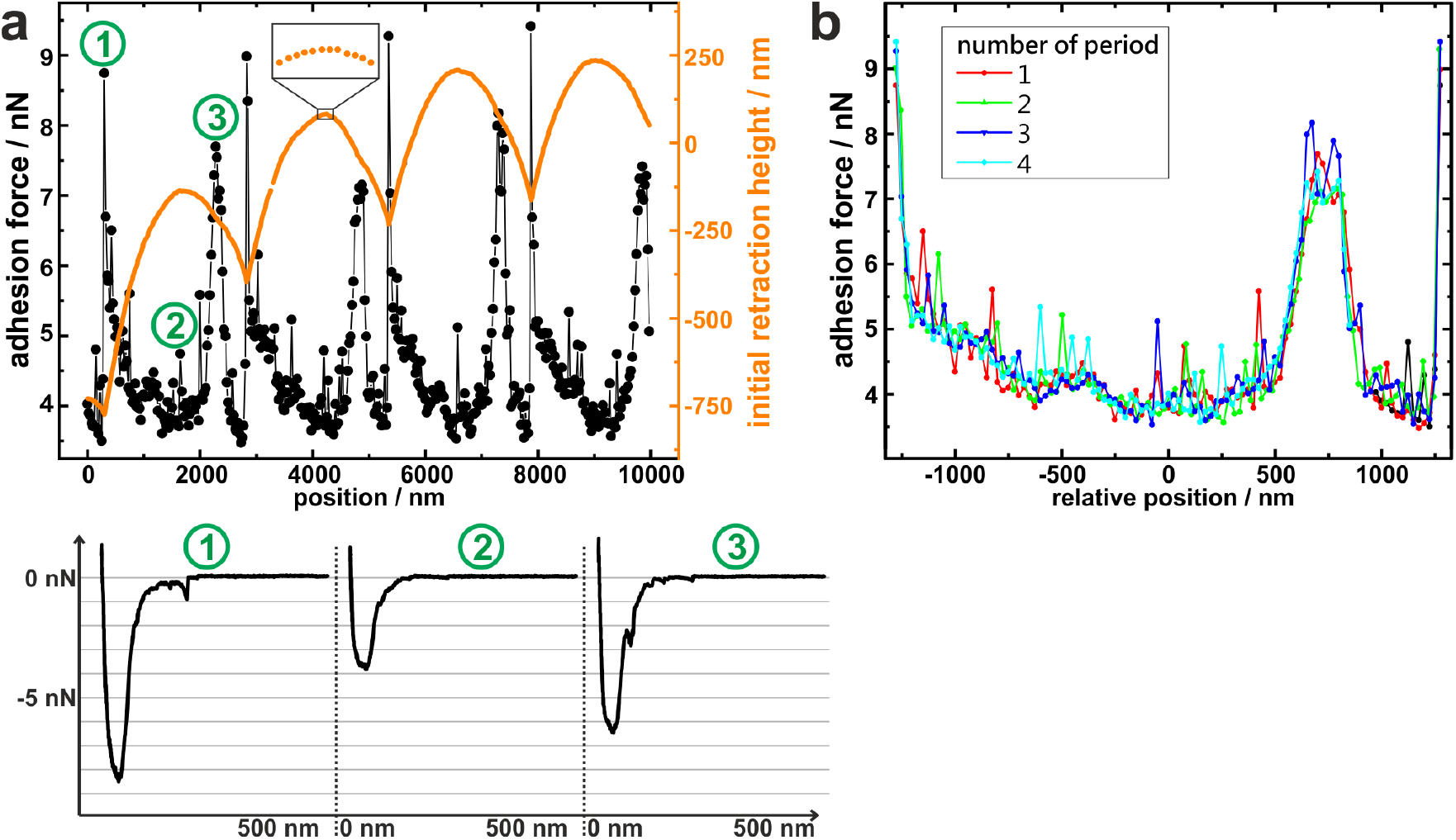
Adhesion force as function of position on the wrinkled surface. Raw data for an exemplary *S. aureus* cell: a) Adhesion force (black dots) and height of the cantilever at which retraction started (“initial retraction height”, orange line) for different positions on the surface. For three positions (1/2: at the minimum/maximum of the surface topography, 3: at an intermediate position with high adhesion), exemplary force-distance curves are shown. The zoom into the orange line highlights the quality of the initial retraction height data. b) Overlay of the adhesion force data from a) for each period of the surface (that was determined using retraction height data) in dependence of the distance from the maximum of each period.

The graph of the initial retraction heights (orange data in Fig. 2a) has a distinct periodicity which reflects the surface topography. Notably, it does not have the same curve form as the AFM scans in Figure 1d. The reason for this is that the AFM tip that scanned the surface had a tip radius of approximately 20 nm while the force-distance curves were recorded with an attached bacterial cell that features a much wider radius (approx. 500 nm). Hence, the cell – in contrast to the tip – cannot exactly follow the surface topography, especially not in the trenches of the surface (For an explanatory sketch, see Fig. S1 in the ESI †). In addition, the AFM has a certain vertical drift that causes a linear offset in the orange data in Figure 2a. Nevertheless, the data reproduce the surface periodicity very well and can be used to extract the position of each force-distance curve in relation to the periodic structures of the substrate.

All recorded force-distance curves (three of which are exemplary shown in Fig. 2a) have a similar cup-like shape, suggesting that a rather high number of cell wall molecules is responsible for adhesion on every position of the wrinkled PDMS^15^. Notably, the recorded adhesion forces show a periodicity with the same wavelength as the initial retraction heights: For example, the graph of the adhesion forces has local maxima at *x* = 200 nm, 2800 nm, 5200 nm, and 7800 nm, each of which nicely corresponds to a minimum in the initial retraction height data.

Next, the recorded adhesion forces were subdivided relative to the periodicity and plotted accordingly, as shown in Figure 2b. In this graph, the recorded adhesion forces inside each period show clearly the same dependence on the surfaces’ topography. This allows us to meaningful average over the results from different periods and construct mean adhesion curves in respect to the surface periodicity (see Fig. 3). These are called adhesion profiles hereinafter and allow us to characterize the adhesion in detail in the next paragraph.

**Fig. 3.**
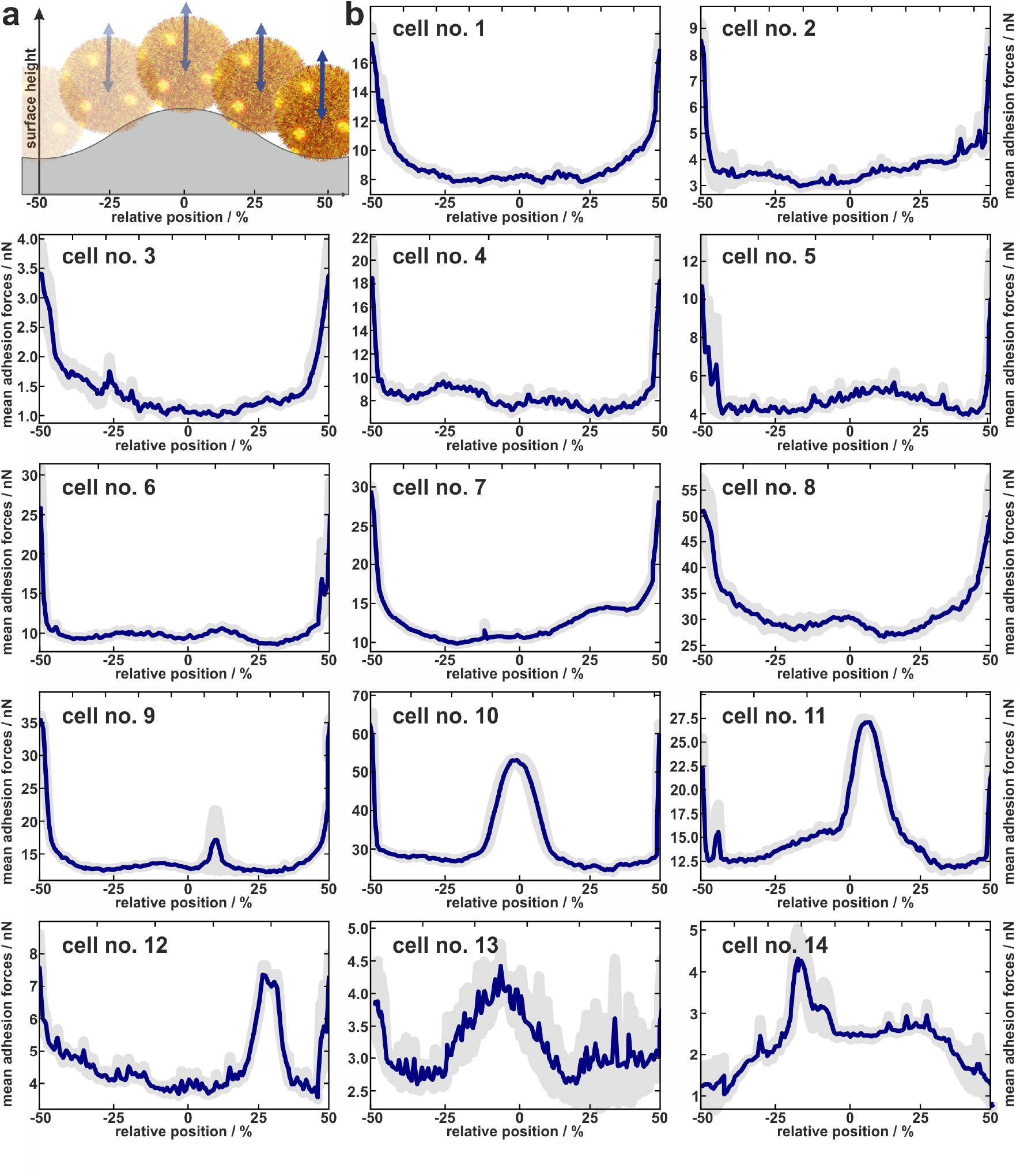
Experimentally determined distribution of adhesion forces of 14 cells. a) About to scale sketch to illustrate how the x-positions fit to the relative position within one period of the surface. b) Mean adhesion forces (and error of the mean as shaded area) of 14 cells averaged over several periods of the wrinkled surface relative to the topographical maximum of the surface periods.

### 1.3 Individual Adhesion Profiles of Experimentally Tested Cells

In total, the adhesion profiles for 14 individual *S. aureus* cells were construed as described in the previous paragraph. The beginning and end of the x-axes refer to the minima of the surface (“trenches”) while the middle corresponds to the local maxima (“hills”) as shown in Figure 3a.

Cells no. 1-12 show comparatively high adhesion forces at the minima of the surface. Between these maxima, the mean adhesion forces are up to a factor of three smaller but feature local maxima that are more or less pronounced depending on the individual cell.

For example, cell no. 1-8 show a bathtub-like adhesion profile with only small local maxima. These profiles are asymmetric around the surfaces’ hill (*F*_*adh*_(*x*)≠*F*_*adh*_(*−x*)), though the surface topography (reflected by the initial retraction heights of 2a) is highly symmetrical. In contrast, cells no. 9-14 feature very pronounced local maxima between the maxima of the curves.

Notably, the existence of local maxima or their relative size does not depend on the measured mean adhesion forces (i.e. the mean value of the measured adhesion forces on every position). In other words, cells with rather low overall adhesive strength can have distinct positions with relatively high adhesion (e. g. cell no. 13), while other cells with a rather strong overall adhesion do not show these positions (e. g. cell no. 6). The cells no. 13 and no. 14, for instance, even do not show the highest adhesion in the minima of the surface. Apparently, the adhesion strength of their highly adhesive positions on the cell wall surpasses the effect of increased contact area-enhanced stronger adhesion in the period’s minima.

To summarize, the measured adhesion forces of the tested cells not only show often distinct local maxima in the surfaces’ minima, but sometimes enhanced adhesion capabilities outside these minima. Furthermore, even for adhesion profiles without additional peaks, i.e. outside the surfaces minima, the profiles are asymmetric within a period. Hence the adhesion capabilities are clearly heterogeneously distributed over the cell envelope. In order to interpret the profiles, it is important to note that we do not necessarily measure the same adhesion forces on the wrinkled surface as we would on a flat substrate: By moving along the surface, not only the location of the bacterial surface area that can contribute to the adhesion force changes, but also its size. Furthermore, since the cantilever with the bacterial cell moves only in a vertical direction, the direction of the cell’s movement relative to the local normal direction of the surface changes for different positions within one period. Since the mechanic properties of the involved macromolecules during elongation under different angles are unknown, it is not straightforward to correct the measured values for this geometric effect and thus directly determine the distribution of adhesins on the cell wall. Therefore, we attempted to disentangle the contributions of the varying bacterial surface area from the mechanical stretching by comparing the experimentally measured data with simulations, as described in the next section.

### 1.4 Disentangling the Origin of the Adhesion Profiles

To disentangle the influence of a varying bacterial surface area from the influence of distinct mechanical stretching of macromolecules, we simulated the bacteria as hard spheres on which adhesive molecules are distributed. Since the adhesion process of *S. aureus* is governed by the collective response of individual macromolecules to stretching^15,16, 34^, whose mechanical properties, e. g. length, and stiffness, are heterogenous and can lead to macroscopically nonlinear behavior in SCFS experiments^34^, we used the model published by Maikranz et al.^15^ and extended it to curved surfaces (see ESI †). Most importantly we included the possibility for an angle-dependent molecule-substratum interaction, which are typically associated with complex protein mixtures. To estimated the bacterial surface area that can interact with the surface, we used a rather simple geometric model, where adhesive molecules are modelled as rods of fixed length that protrude outward the bacterial cell wall that is modelled as a hard sphere. After the cell is brought into tangential contact at position x above the surface, the relative adhesive strength is calculated from the molecules intersecting with the surface (each molecule might also have a different adhesiveness, for details, see Materials and Methods Section and ESI †). In essence, the geometric model describes, in the limit of many uniform distributed macromolecules, the fraction of the bacterial surface area that is able to contribute to the adhesion (see Fig. S2 for how this depends on the number of macromolecules). Hence, the geometric model does not provide the correct force scale but relative values.

#### 1.4.1 Patches of Molecules

Before we disentangle the origin of the underlying adhesion profiles we briefly present how peaked adhesion capabilities outside the surfaces trenches can be created. To this end, we systematically placed a single patch of variable size on the bacterium and obtained the adhesion profile in the geometric model (see Fig. S3 and S4). These single patches reproduced isolated peaks at the surfaces maximum when placed at the bottom of the bacterium. While this results is expected, interestingly, for no other position along the surface isolated peaks could be obtained by a single patch. These rather produce shoulders or parabola-like profiles when the patch does not contribute to the adhesion at all. When several patches were distributed either independently, or with a certain distance to each other, we obtained asymmetric parabola-like profiles (see Fig. 4. The asymmetric parabola-like profiles are obtained for a rather large number (up to 30) of independent patches with diameters of about 50 nm, while the peaked profiles are caused by distinct patches (about 5-6 patches, some of which have a distance of at least 850 nm to neighbouring patches) with a larger diameter of about 250 nm. Inside the patches, the number of molecules or their individual adhesive strength are enhanced (by about a factor of 15, see also Fig. S5). Now that we clarified how peaked adhesion capabilities at the surfaces maximum can be created, we disentangled how homogeneously distributed molecules create the underlying profile.

**Fig. 4.**
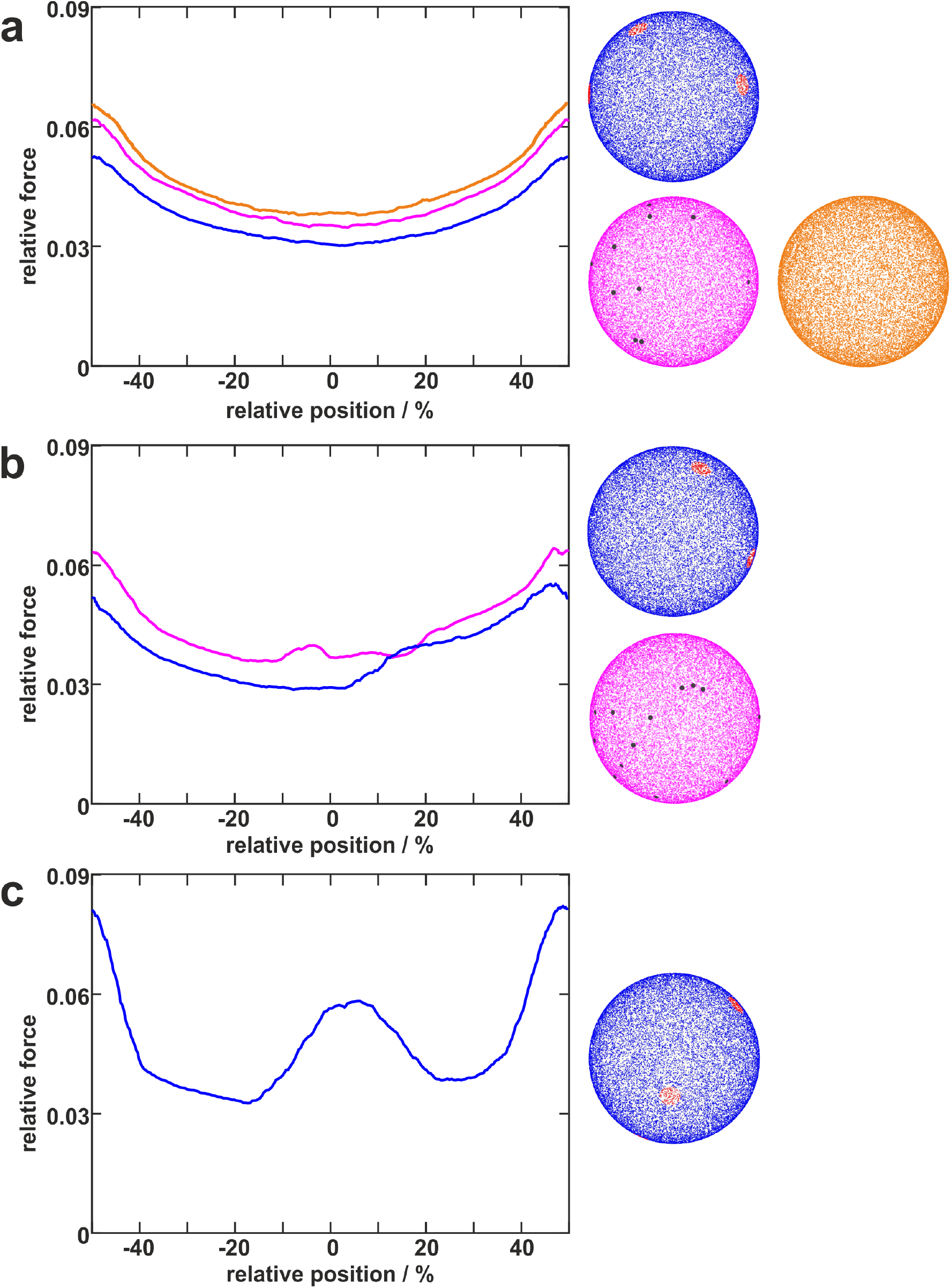
Distribution of adhesive molecules and corresponding simulation results. Examples of simulated cells with differently distributed adhesive molecules (right) and corresponding adhesion force profiles (left). The blue cells with distinct large patches (red) with a certain distance to each other can reproduce several types of profiles (blue lines in a, b, c). In contrast, the pink cells having independently distributed small patches (darkgrey) can only reproduce rather smooth curves (pink line in a) or curves with only small “humps” (pink lines in b). The orange cell with homogeneously distributed molecules can only reproduce smooth cup-shaped profiles (orange line in a).

#### 1.4.2 Homogeneously distributed molecules

Now, we simulated the adhesion profiles originating from homogeneously distributed molecules. To this end, we compared the simulations with a mean adhesion profile constructed from the experimentally obtained profiles which displayed only in the surfaces trenches increased adhesion capabilities (cells 1-8). Since the geometric model only provides relative values, we normalize all adhesion forces by the maximal value. Note, that this also reduces the influence of the rod length onto the adhesion profile in the geometric model. Furthermore, the model considering the mechanical stretching is able to reproduce the experimentally observed force scales (see Fig. S6).

As in the experiments all models showed the maximal obtained adhesion in the surfaces trench, which then decreased towards the surfaces’ maximum. Note, however, that for varying properties of the macromolecules the maximal adhesion force is not necessarily realized (see Fig. S6). A comparison of the geometric model and the thermally fluctuating molecules in the absence of angle-dependent interactions show that although the geometric model produced smoother curves, the overall shapes and relative magnitudes of the profiles obtained from both models were the same (see Fig. 5). Thus, both models displayed rather parabola-like profiles instead of the experimentally observed bathtub. However, if we consider angle-dependent interactions, a bathtub-like adhesion profile was recovered. The influence of the angle-dependence is emphasised by rescaling into the adhesion force in the geometric model by the magnitude of the local surface normal (see Fig. 5). This leads to a smaller extend of the plateau in the adhesion profile. Hence the interaction of individual macromolecules with the local surface potential is expressed in the length of the plateau. However, none of the considered models reproduced the magnitude of the reduction in adhesion forces to about 50%.

**Fig. 5.**
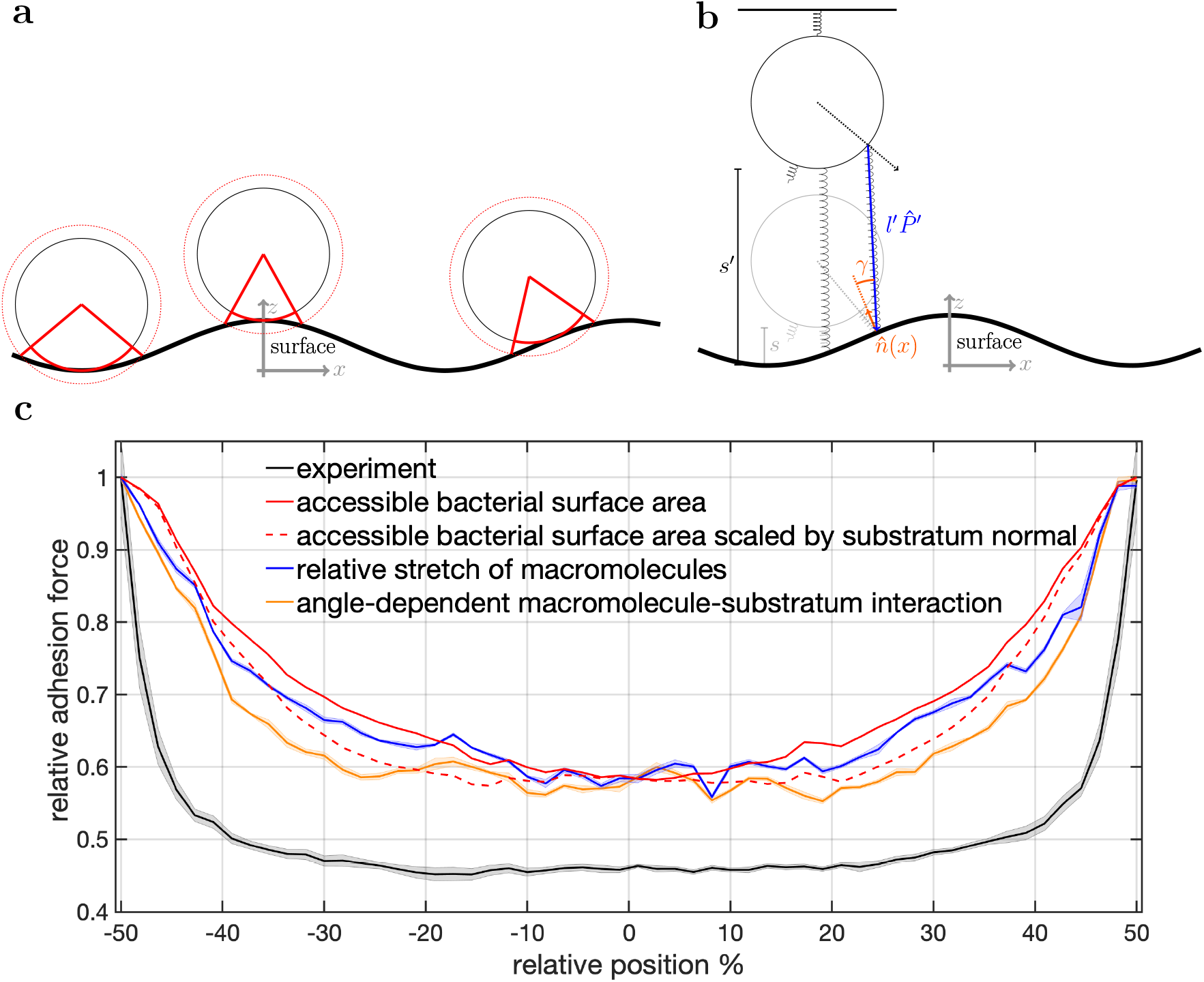
Model schematics and resulting mean adhesion profiles. a) Illustration of the varying bacterial surface area that is able to interact with the surface when a certain interaction range indicated by the dashed line is assumed. b) Illustration of the stretch tethered macromolecules experience when the bacterium is retracted and an illustration of the possible angle-dependent macromolecule-substratum interaction. c) Resulting adhesion profiles normalized by the maximal adhesion force. For the simulation results we averaged over 5 repetitions of 5 distinct bacteria, while for the experimental mean adhesion profile we averaged cells 1-8 in Figure 2b. Note that only angle-dependent macromolecule-substratum interaction reproduce a constant profile along large distances of the surface.

To understand the reduced magnitude, we tracked in the model not only the total force experienced by the bacterium but also its spatial origin (see Fig. S8). This analysis (for a discussion of the results see the ESI †) revealed that inside the surfaces minimum mostly molecules pointing perpendicular to the surfaces periodicity contribute to the adhesion force. While outside the surfaces minimum the molecules pointing along the trench contribute probably the most to the adhesion. A naive attempt to exploit this feature by introducing a cut-off angle for the molecule-substratum interaction lead for small cut-offs to a marginal extension of the adhesion plateau but not to a reduction in adhesion force (see Fig. S8). In fact, larger cut-offs led only away from the surfaces maximum to a reduction in adhesion forces such that no bathtub-like profile was recovered. Hence, more complicated molecule-substratum interactions should be considered. However, since our model considers the bacterium as a rigid sphere, the influence of deformations and elliptical shape were not considered. While these effects, as well as neglected substratum deformations, are considered to be secondary factors, a true evaluation of the angle-dependence requires the repetition of SCFS experiments, whereby the bacteria are retracted at an angle to the surface. This, however, requires a specialised experimental set-up and is beyond the scope of this work.

## 2 Conclusions

We investigated the adhesion capability of *S. aureus* cells to a periodically structured surface by single-cell force spectroscopy to measure how the strength of adhesion depends on the position relative to a structured surface. We found that the adhesion of bacteria is not only cell-specific (as shown before (15, 17)), but also depends on the position on the cell envelope. Simulations reproducing the experimental results revealed the importance of angle-dependent interactions, and gave information about the distribution of the adhesion capability on the cell wall: Our data show a large reduction of adhesion forces outside of the surfaces minimum but also that *S. aureus* cells can have highly adhesive patches. Depending on the probed cell, these patches have different properties: While the experimental results for some cells suggest a rather high number (up to 30) of independent patches with diameters of about 50 nm, other cells must have fewer distinct patches (about 5-6 patches, some of which have a distance of at least 850 nm to neighbouring patches) with a larger diameter of about 250 nm.

Hence, our results for coccal-shaped Gram-positive *S. aureus* cells nicely complement the patchy colloid model of adhesion for rod-shaped Gram-negative *E. coli* cells by Vissers et al.^30^. Their experiments show that *E. coli* cells have distinct patches on their surface and that the number of these patches defines adhesive strength of a cell; if no patches exist, a cell hardly adheres. However, our results – together with former studies – lead to a slightly different notion for *S. aureus* cells: Since the force-distance curves on all positions of the surface look similar, namely cup-shaped, *S. aureus* cells seem to have many adhesive molecules at almost every position of the cell wall, but the strength of adhesion has maxima at certain locations^15^.

At these points, not necessarily the number of molecules is maximal, but rather their individual properties lead to maximum adhesive strength^17^. Although we do not determine the origin of the adhesive patches, the angle dependence of the interaction suggests molecules with complex 3d structures, like, proteins as the source. Furthermore, the simulated patchy spheres (see Fig. 4) are quite similar to the electron micrographs showing the distribution of protein A and Clumping factor A in the publication of Harris et al.^21^. Other candidates for the origin of the adhesive patches might be Cna and/or FnbpA, since both are multifunctional adhesins and cluster in nanometer-sized domains on the *S. aureus* cell wall^26,35^. However, we cannot answer the question, whether the adhesive patches are “hot spots” where many adhesins occur together, whether there are several clusters, each containing only one type of adhesin, or whether the combination of different adhesive molecules with certain mechanical properties renders a given position at the cell wall highly adhesive.

Moreover, we cannot resolve if cells that do not show very distinct maxima in the adhesion profiles do not have any patches, or if – by chance – none of the patches come in contact to the surface. Along this line, it might be possible that only one half of the cell, for example the part that was newly synthesized during cell division, has patches of high adhesion capability^36,37^. This is an exciting subject for further studies, in which adhesion measurements on structured surfaces could be combined with fluorescent labelling techniques^38^. In that way, it will be possible to correlate the prevalence of certain proteins and/or former division planes with the adhesion capability of the investigated cells. Alternatively or in addition to this, extracellular vesicles formed and temporary retained on the *S. aureus* cell surface might contribute to this phenomenon^39,40^.

Our findings have consequences for science and material development: In future experiments and especially when designing models for simulations, the cells should not be regarded as rather uniform colloids, but as objects with heterogeneous surface properties. Finally, these differences in adhesive properties should be considered in the design of new antibacterial materials for the reduction of infections. In particular, the large reduction of adhesion forces outside the surfaces trenches, caused by an angle-dependent substratum interaction, could be exploited to reduce adhesion.

## 3 Material and Methods

### 3.1 Production of the Wrinkled Surfaces

PDMS was prepared by mixing the pre-polymer and curing agent of a Dow Corning Sylgard 184 PDMS Kit in 5:1 ratio, curing it at RT for 24 h followed by a thermal treatment of 4 h at 80 ^*°*^*C* under ambient conditions. Slabs of 4.5 cm *×* 1.0 cm were cut out, cleaned with Milli-Q water and dried with nitrogen. The slabs were clamped in a custom-made stretching-device and strained uniaxially to 5–10 % of their initial length. Afterwards the slabs were placed in a low-pressure RF-plasma chamber and treated for 120–300 s with a H2-plasma at 800 W. Eventually the prestrain is released, revealing opaque colored wrinkles on the PDMS topside^31^.

### 3.2 Bacterial Cultures

*S. aureus* cells, strain SA113, from a deep-frozen stock culture were plated on blood agar for one day and a fresh plate was used no longer than a week. The day before the experiments, one colony from the plate was transferred into 5 ml of tryptic soy broth (TSB) and cultured for 16 h at 37 ^*°*^*C* under agitation (150 rpm). To get cells in exponential growth phase, at the day of the experiments, 40 *μ*l of the overnight culture were transferred into 4 ml of fresh TSB and cultured for 2.5 h at 37 ^*°*^*C* under agitation (150 rpm). From this final culture, 1 ml was washed three times with sterile phosphate buffered saline (PBS) at an acceleration of 17,000 g. The cells in PBS were stored at 4 ^*°*^*C* and used no longer than 6 h.

### 3.3 Single-Cell Force Spectroscopy

As described in the publication of Thewes et al., using a micromanipulator (Narishige Group, Tokyo, Japan), single bacterial cells were immobilized on tipless cantilevers (MLCT-0-F with nominal spring constants of 0.03 N/m from Bruker, Santa Barbara), which were beforehand coated with polydopamine^41^. With these bacterial probes, single-cell force spectroscopy measurements were performed using a Bioscope Catalyst (Bruker) at room temperature in PBS (pH 7.3, ionic strength 0.1728 mol/l). Force-distance curves were performed with a ramp size of 800 nm and a velocity of 800 nm/s. The force trigger, i.e. the maximal force with which the cell is pressed onto the substrate prior to immediate retraction, was set to 300 pN. With every cell, some hundreds (between 400 and 500) of consecutive curves were recorded in a straight line with a constant lateral distance (of 20 nm, 25 nm, or 30 nm; called x-offset hereinafter) between consecutive curves on one of the three PDMS samples. Hence, force measurements on 4–5 equivalent positions in different periods were recorded. No systematic change in the adhesion behavior, such as a decreasing adhesive strength due to cell fatigue, could be observed even after 500 curves. The direction of this straight line was perpendicular to the trenches in the wrinkled PDMS samples with a deviation of less than 1^*°*^. For every probed cell, the parameters of the experiment (number of curves, x-offset, underlaying PDMS sample) are given in Table S1 in the Supporting Information.

### 3.4 Analysis

From every recorded force-distance curve, a baseline was first subtracted, and the adhesion force was determined as the minimum force that occurred during retraction of the cantilever. In addition, the z-position of the instrument’s height sensor at the beginning of the retraction was recorded, and also corrected for a linear baseline shift caused by a drift of the AFM piezo. The adhesion forces and the positions where the retraction of the cantilever started were plotted against the corresponding x-offset and the periodicity was determined automatically as follows: The curves of the initial retraction heights were searched for peaks in negative direction (denoting the valleys of the surface). The positions of these peaks were used to divide the calculated adhesion forces into sections that correspond to the different periods of the surface. Since the wavelength of the periodicity can locally vary and since it does not necessarily fit a multiple of the x-offset, the data for each period were slightly shifted in x-direction, so that each period has the same size.

### 3.5 Simulations

To obtain an estimate of the distribution of adhesive molecules on the bacterial cell wall, we used two types of models:

#### Monte Carlo Model – Thermally Fluctuating Macromolecules

Since the used substrate surface is curved, geometric constraints of the surface-sphere geometry non-uniformly affect the interactions of the macromolecules along the substrate. Therefore, we extended the model of Maikranz et al.^15^, in which the bacterium is modelled as a hard sphere, decorated with thermally fluctuating macromolecules. For the mechanical response a WLC polymer model with probabilistic parameters is used where the macromolecule is stretched when it is bound to the substrate. In the extension, we consider thermal length fluctuations and acting forces along the respective normals of each macromolecule. Most importantly we include the possibility for an angle depend molecule-substratum interaction (for model details, see ESI †).

#### Geometric Model

For the bacterium, a hard sphere with a radius of 500 nm was used. Adhesive molecules in the cell envelope, were modelled as rods of constant lengths protruding from the surface of the cell outwards in normal direction. The model provides the relative adhesion force of the cell at point x along the sinusoidal surface by a weighted count of all rods intersecting the sine surface in relation to the total number of rods (for details, see ESI †).

#### Distribution of Adhesive Capabilities

We investigated homogeneous as well as patchy distributions of adhesive capabilities. A random, homogeneous distribution of rods on the cell surface was realized by placing rods with identical weight randomly on the sphere (independently and uniformly distributed). We either uniformly distribute 50.000 molecules and compare the results of the thermally fluctuating macromolecules with the geometric model. Or, to obtain complete spatial randomness, the number of rods follow a Poisson distribution^42^. Adhesive patches have been produced by placing clusters of fixed radial extension (spherical caps) onto a homogeneous distribution. Inside these clusters the adhesive strength was increased by either placing additional rods inside the cluster or by giving all rods inside a cluster a larger adhesive weight. These clusters have been realized in two different ways: i) Clusters via random position: A Poisson distributed number of spherical caps with constant radii (125 nm) were placed at random positions. Note that different clusters might overlap. ii) Clusters via random sequential adsorption (RSA)^43^: In each adsorption step, a spherical cap with constant radius (125 nm) and a constant “radius of repulsion” (850 nm) was randomly positioned on the spheres surface. The position of the following spherical caps was only accepted if their “radii of repulsion” did not overlap with previously placed caps. The RSA process was stopped after 1000 runs, i. e. when with a high probability no additional spherical caps could be added.

## Acknowledgements

The authors thank the German Research Foundation (DFG) for funding within the context of the Collaborative Research Center SFB 1027 (projects B1 and B2). C. S. acknowledges funding from the DFG the project JA 905/6. M. A. K. acknowledges funding by the Princeton University Innovation Fund for New Ideas in the Natural Sciences and support by tby the Deutsche Forschungsge-meinschaft (DFG, German Research Foundation) through the SPP 2265, under grants numbers ME 1361/16-1, WI 5527/1-1, and LO 418/25-1, as well as by the Volkswagenstiftung via the Experiment Project “Finite Projective Geometry.” A. F. and B. G. acknowledge funding from the DFG project number FE 600/20-1. K. J. acknowledges funding by the Deutsche Forschungsgemeinschaft (DFG, German Research Foundation) priority program SPP 2265 under grant number JA 905/8-1 and DFG large instrument funding under grant number INST 256/542-1 FUGG (project number 449375068) as well as funding within the Max Planck School Matter to Life supported by the German Federal Ministry of Education and Research (BMBF) in collaboration with the Max Planck Society.

## Footnotes

§ We acknowledge that at low densities it is difficult to meaningfully define heterogeneously distributed.

