## Additional Figs. S1 - S11 and Table T1. for "The adhesion capability of *Staphylococcus aureus* cells is heterogeneously distributed over the cell envelope"

Cite this: DOI: 00.0000/xxxxxxxxxx

### Electronic Supplementary Information for 'The adhesion capability of *Staphylococcus aureus* cells is heterogeneously distributed over the cell envelope'

Christian Spengler[a,‡], Erik Maikranz[b,‡], Bernhard Glatz[c], Michael Andreas Klatt[d,*], Hannah Heintz[a], Markus Bischoff[e], Ludger Santen[b], Andreas Fery[f] and Karin Jacobs[a,¶]

#### 1 Supplementary Figures

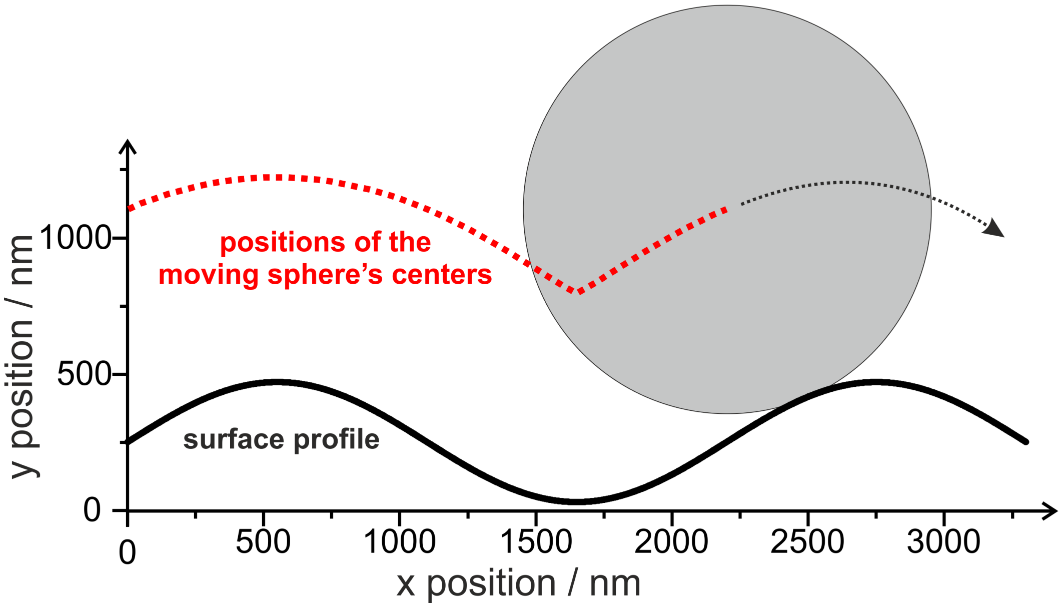


**Fig. S1 "Scanning" with a cell.** The image sketches the expected scan line (red dots) recorded with the sketched sphere on the shown sinusoidal surface (black line). At $x = 1600$ nm, the scan line features a "kink" (exaggerate in the schematic) that is not present at the surface. The red curve is in nice accordance with the data from the AFM height sensor shown in Figure 2a. Note that we were not actually scanning with a bacterium, rather every red dot symbolizes the initial retraction height.

[a]Experimental Physics, Saarland University, Center for Biophysics, 66123 Saarbrücken, Germany
[b]Theoretical Physics, Saarland University, Center for Biophysics, 66123 Saarbrücken, Germany
[c]Institute of Physical Chemistry and Physics of Polymers, Leibniz Institute of Polymer Research, 01069 Dresden, Germany
[d]Department of Physics, Princeton University, Jadwin Hall, Princeton, NJ 08544-0001, USA
[e]Insitute of Medical Microbiology and Hygiene, Saarland University, Center for Biophysics, 66421 Homburg/Saar, Germany
[f]Physical Chemistry of Polymer Materials, Technical University Dresden, 01062 Dresden, Germany
† Electronic Supplementary Information (ESI) available: See DOI: 00.0000/00000000.
‡ These authors contributed equally to this work.
* Present address: Institut für Theoretische Physik II: Weiche Materie, Heinrich-Heine-Universität Düsseldorf, 40225 Düsseldorf, Germany; Institute of Theoretical Physics, Friedrich-Alexander-Universität Erlangen-Nürnberg (FAU), 91058 Erlangen, Germany, and Experimental Physics, Saarland University, 66123 Saarbrücken, Germany
¶ Max Planck School Matter to Life, 69120 Heidelberg, Germany

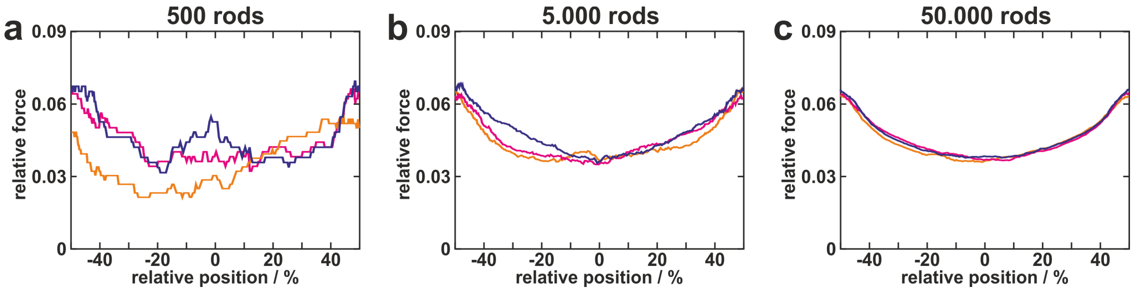


**Fig. S2 Influence of the number of adhesive molecules.** The graphs show adhesion profiles of three simulated cells each with different numbers (a: 500, b: 5.000, c: 50.000) of homogeneously distributed adhesive molecules in the geometric model.

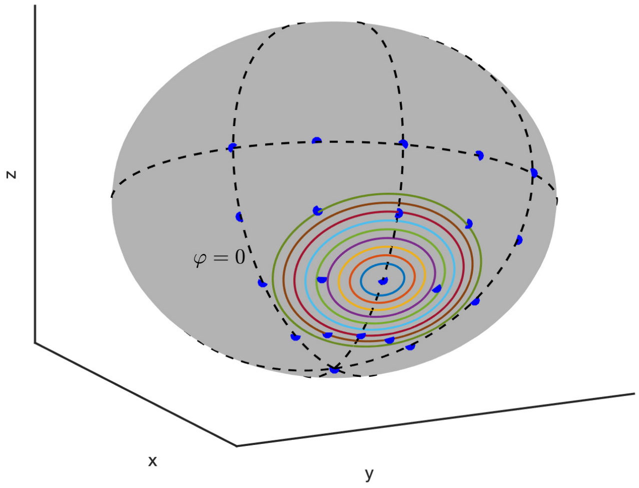


**Fig. S3 Illustration of the patches positions in Figure S4.** The radii of the patch are up to scale and match the colours in Figure S4. Note that in the simulations $\varphi = 0$ aligns with movement parallel to the surfaces' periodicity.

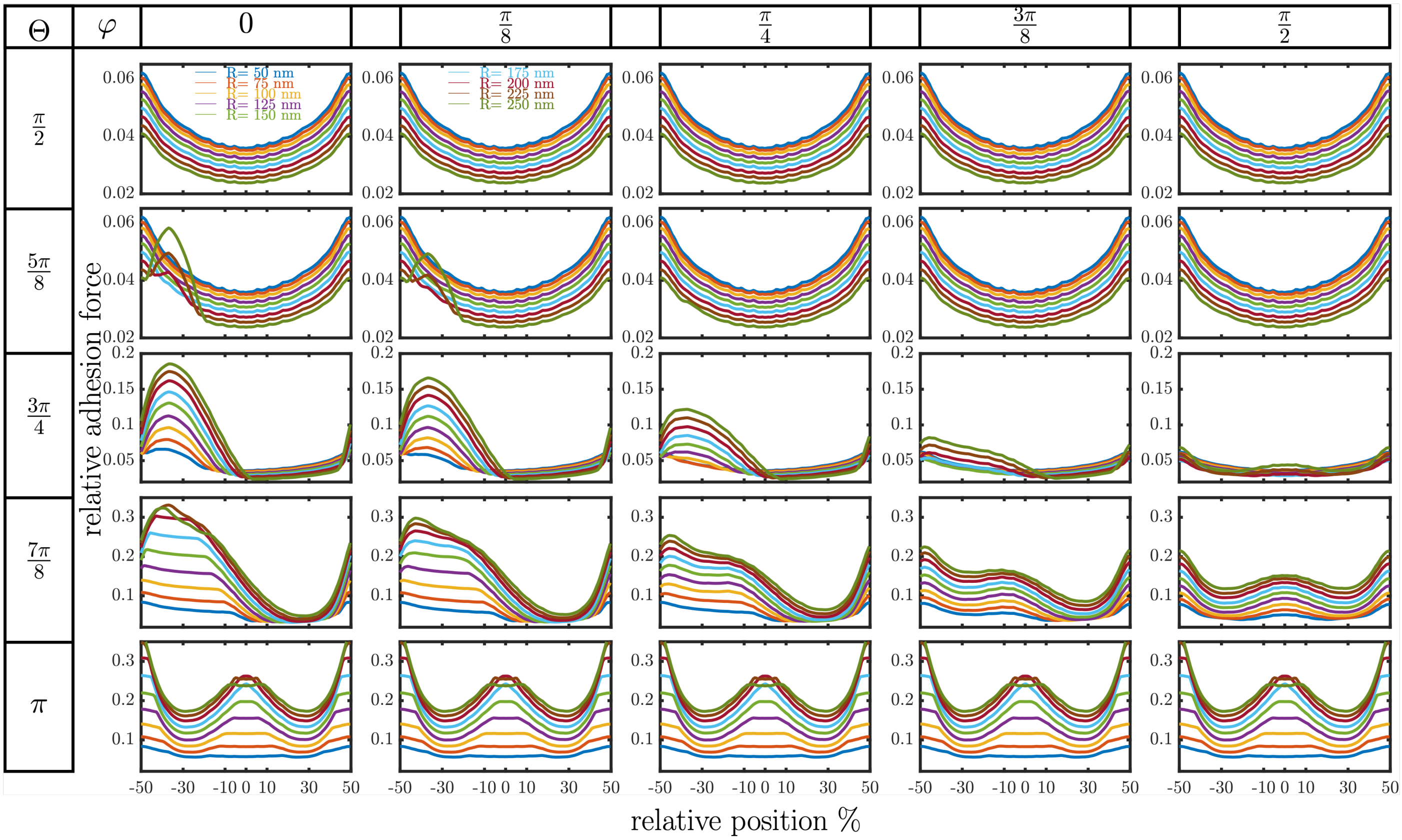


**Fig. S4 Adhesion profiles of the single patches positioned as shown in Figure S3.** Adhesion profiles in the modified geometric model with a single patch of radius R and ten-fold increased adhesiveness positioned at position $(\Theta, \varphi)$, whereby $\Theta, \varphi$ are the usual polar and azimuthal angle in spherical polar coordinates. Note that in the simulations $\varphi = 0$ aligns with movement parallel to the surfaces' periodicity.

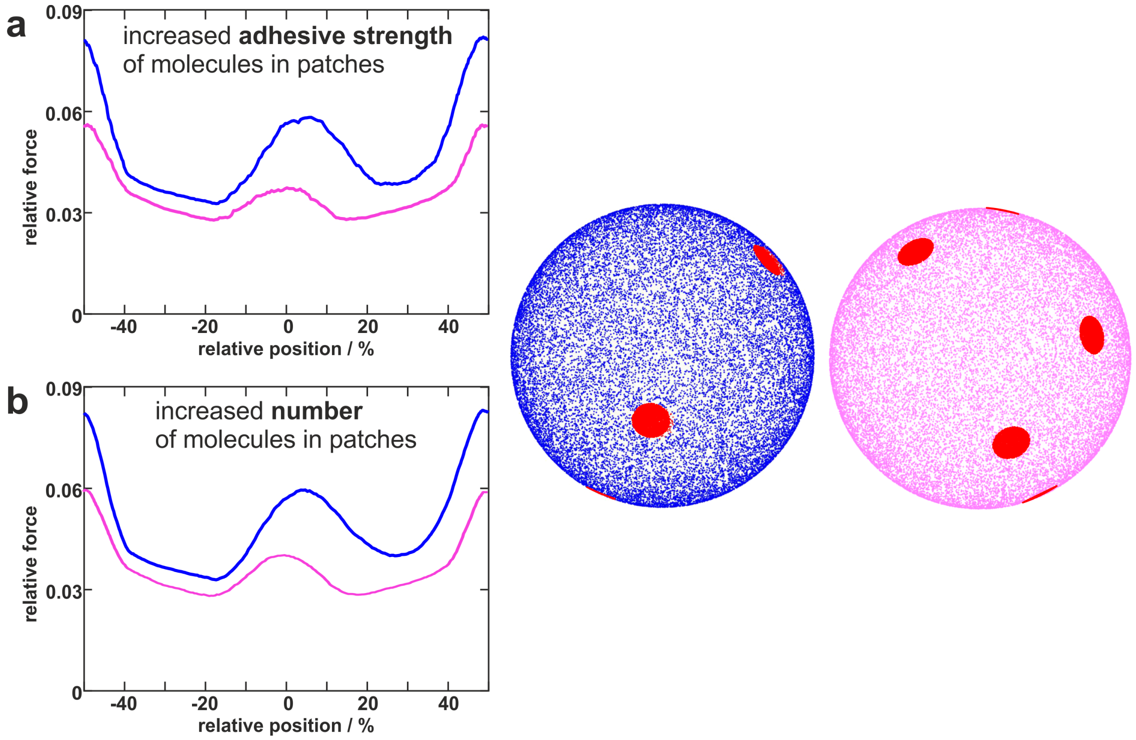


**Fig. S5 Simulations with increased adhesive strength vs. increased molecule density.** The figure shows two curves each for the adhesion profile of two cells (blue and pink) where the molecules in the patches (indicated in red) have a 15-fold increased adhesive strength (a) or 15-fold increased density (b).

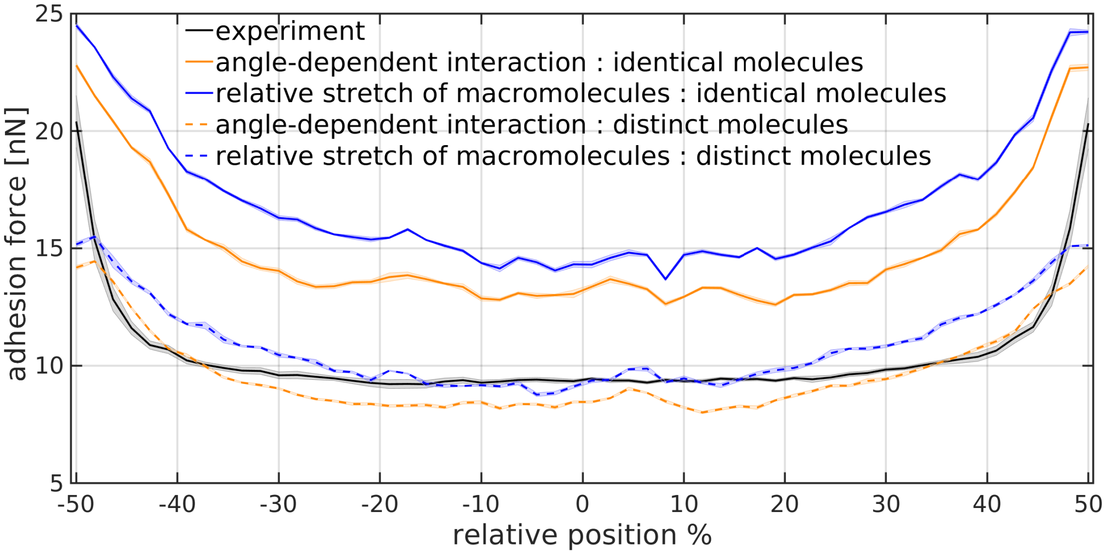


**Fig. S6 Comparing the force scale.** Comparison of the Monte Carlo simulations with the experimentally obtained mean profile averaged for cells 1-8. In the simulations 50.000 molecules are uniformly distributed, over the entire bacterium. Binding energy $V = 20k_BT$. Continuous line (shaded area is SEM of five repeated simulations) are the adhesion profiles for identical molecules with contour length of 45 nm and a Kuhn length of 0.4 nm. The dashed lines represented adhesion profiles of uniformly distributed molecules whose Kuhn length are uniformly distributed in 0.2 -1 nm, while the contour length is Weibull distributed with form parameter 1.5 and scale parameter 125 nm. Blue and orange curves represent the same bacterium, and thus just the considered molecule-substratum interaction is changed. Note that when the properties are not uniform the maximal adhesion force is not directly obtained in the surfaces minimum.

#### 2 Spatial Resolution of the Varying Bacterial Surface Area Along the Surface

As a first step to resolve the varying contribution of the bacterial surface area, we computed for every point along the surface the contact point of the bacterium (see Figure S7, inset) as described in section S4. Note that while moving from the surfaces' minimum to its maximum the same contact point is reached twice (see Figure S7). The molecules binding to the substrate are expected to be symmetrical distributed around the contact point. Hence, in the following we will use the contact point to define the origin of a coordinate system, which changes along the surface (see Figure S8). The new coordinate system is obtained by turning the global z-axis onto the contact point (see Figure S8. This is the same transformation as used in section S4 and there explained in detail). In this new coordinate system we obtain as indicated in Figure S8 a new azimuth and polar angle.

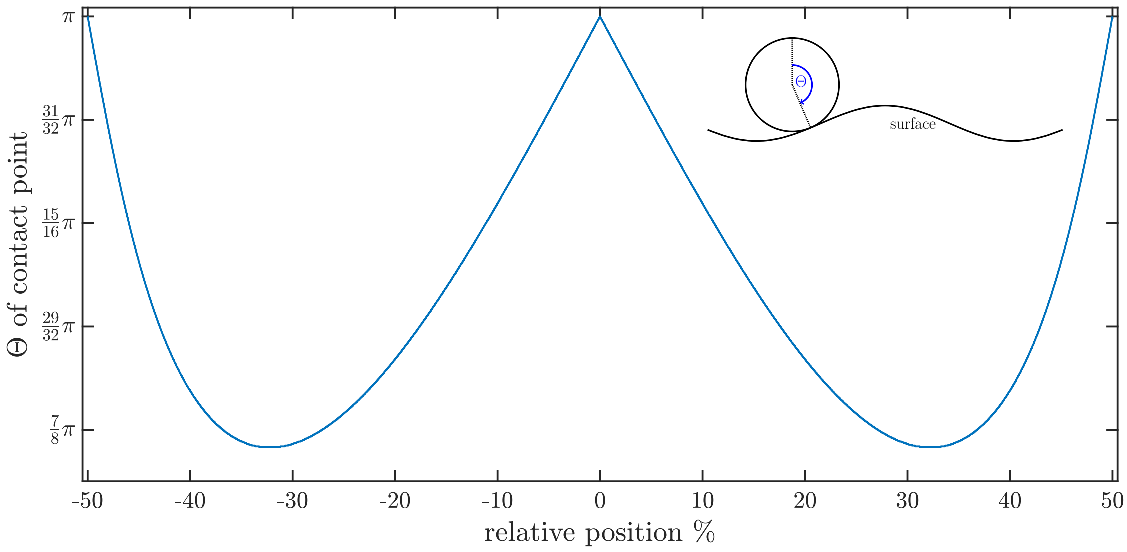


**Fig. S7** Polar angle of the contact point as depicted in the inset. The angle is computed as described in section S5.

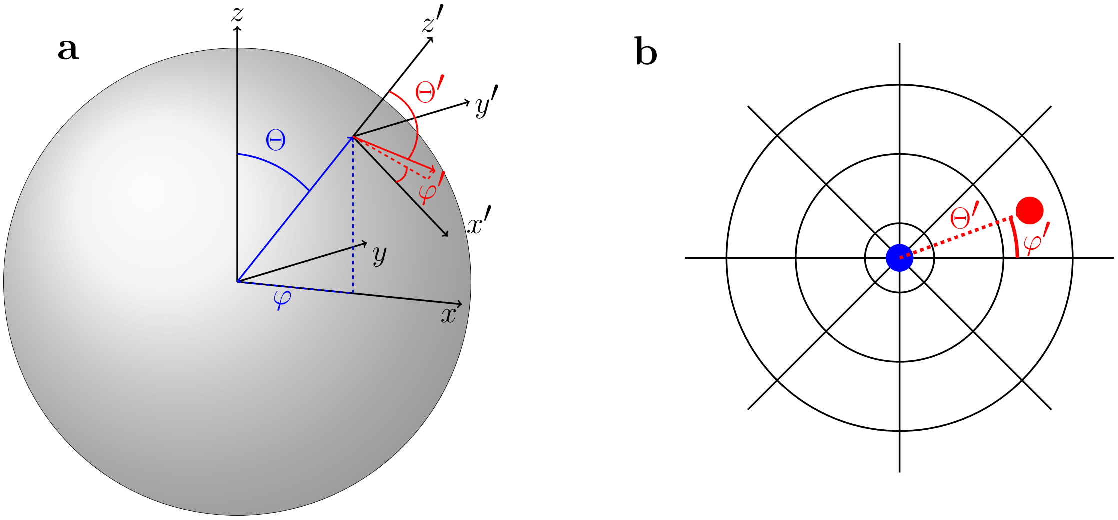


**Fig. S8 Illustration of the coordinate change induced by the changing contact point along the surface.** The coordinate system changes along the x-coordinate, such that the z-axis matches the surface normal of the contact point. The required turning angle is depicted in Figure S7. In the new coordinate system, the radial coordinate is the polar angle while the angular coordinate is the azimuthal angle in the new coordinate system. Therefore, the radial coordinate depicts the geodesic distance to the contact point, whereby the lines of constant angle are 100 nm apart.

To represent how different parts of the bacterium come into contact, we use a polar plot with the contact point as the origin. In this plot the angular coordinate depicts the azimuthal angle in the new coordinate system, while the radial coordinate depicts the new polar angle and hence the geodesic distance on the sphere to the contact point (see Figure S8). Furthermore, we only depict molecules when they contribute to the adhesion force, i.e. $\langle$ force $\rangle = 0$. Hence the white space is still covered by molecules but they don't contribute to the adhesion force. In accordance with the previous results, the bacterial surface area contributing to the adhesion force is, for the thermally fluctuating molecules, always smaller than in the geometric model (compare purple which is the contribution in the geometric model to other colours in Figure S9). In particular, we observe that in the plots the contributing areas are elliptical around the contact point. These are, however not ellipses in real space. Since this polar plot uses polar and azimuthal angles on a sphere it describes a curved patch. Nevertheless, a smaller area in the polar plot indicates a smaller bacterial surface area.

The reduced area can be attributed to two effects. First, the effective length of the fluctuating macromolecules is much smaller than the interaction radius considered in the geometric model. Second, the geometric model considers the interacting molecules at the point of contact, while the adhesion force for thermally fluctuating molecules is obtained at a certain distance to the surface. And since some molecules detached during retraction, not all initially bound molecules contribute to the adhesion force. This is particularly well highlighted by comparing both interaction potentials (compare Figure S9 a and b). For angular interactions, the area contributing to the force is not simply connected, but close to the contact point, the molecules are mostly detached. This is caused by the effectively smaller potential depth experienced by most of the molecules, which leads to earlier detachment. Interestingly, as can be observed by the relative stretch $\frac{l-l_0}{L}$ , the molecules are stretched symmetrically around the contact point. Increasing the geodesic distance (radial coordinate in Figure S9) the molecules are stretched more until they are detached (white space in Figure S9). Furthermore, all molecules are mostly aligned along the global z-axis such that the force experienced by the molecules is mostly measured by the cantilever (geometric factor in Figure S9).

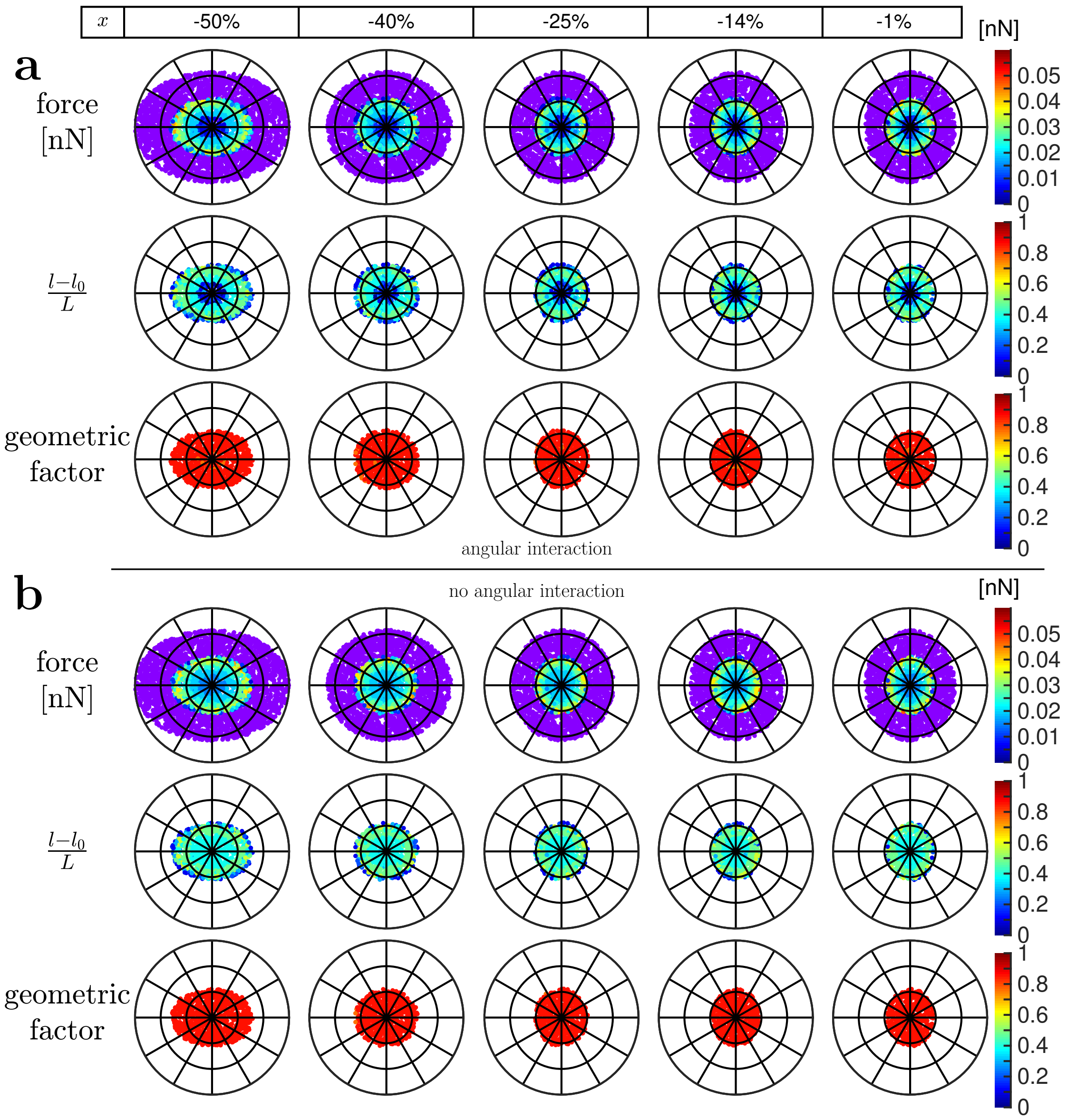

**Fig. S9 Mapping the force contribution on the bacterium's surface.** Molecule properties contributing to the adhesion force for the model including (a) angular interactions and (b) no angular interactions. The coordinate system changes along the x-coordinate, such that the z-axis matches the surface normal of the contact point. The required turning angle is depicted in Figure S7. The radial coordinate is the polar angle while the angular coordinate is the azimuthal angle in the new coordinate system. Therefore, the radial coordinate depicts the geodesic distance to the contact point, whereby the lines of constant angle are 100 nm apart. Purple depicts the molecules which contributed to the force according to the geometric model. Molecules are only plotted when they contribute to the adhesion force, i.e. $\langle$ force $\rangle = 0$. The geometric factor is $P_z$ with respect to the global coordinate system and measures how much of the force experienced by the molecules is transmitted onto the cantilever. All 50.000 molecules have a contour length of 45 nm, a Kuhn length of 0.4 nm and the data is averaged over five retractions.

As I already mentioned the size of the interacting area changes along the surface. However, Figure S9 shows also that the ellipses major axes turn when moving along the surface. Since the local coordinate system turns along the surface caution is warranted in the interpretation. However, the first and last columns, as well as the second and third have approximately the same contact point. Hence, comparing for these the orientation of the ellipses is straightforward. This shows, that in the minimum of the surfaces mostly the molecules pointing along the periodicity of the surface contribute to the attachment. When the bacterium moves out of the minimum of the surface the patch turns. Now the molecules pointing along the trench contribute to the adhesion force.
While we did not track the molecule-substratum angle explicitly, the stored information allowed us to calculate it. Hence we can investigate the influence of an prescribed cut-off angle by computing a set of artificial retraction curves as follows. For each molecule we computed the angle $\gamma$ between the surface normal and the molecule, and if this angle was larger than a prescribed cut-off the molecule is considered as unbound. Thus we obtain a new retraction curve from which the adhesion force can be extracted. As Figure S10 shows a cut-off of 0.5 is identical to no cut-off and a reduction in adhesion-force is not obtained at the surfaces maximum.

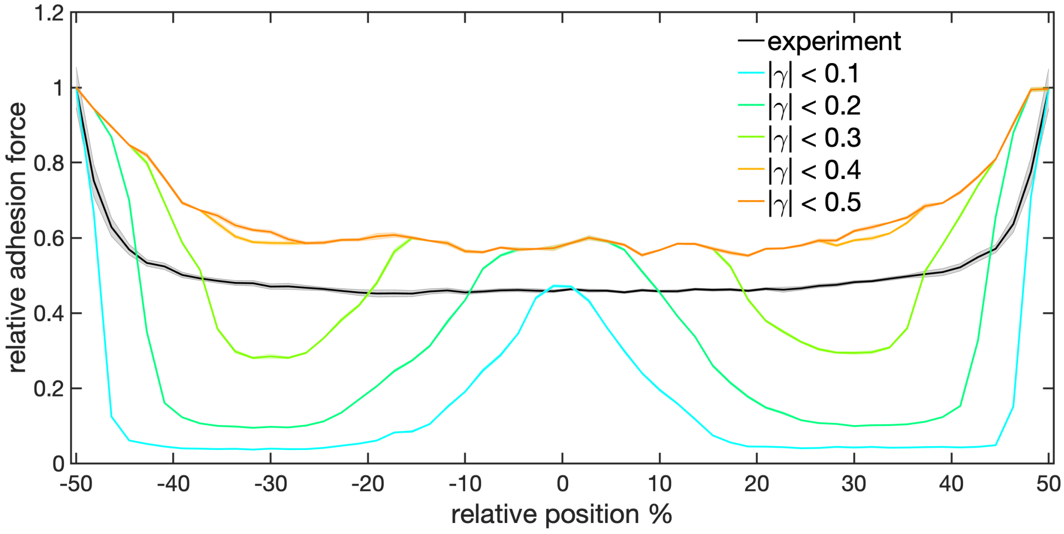


**Fig. S10 Influence of an angle cut-off onto the adhesion profiles.** Cut-off 0.5 is identical to no cut-off.

#### 3 Detailed Description of the Models

**Monte Carlo Model – Thermally Fluctuating Macromolecules** We determine the position $\vec{r}$ of the macromolecules on the bacterium by the usual spherical coordinates $(\Theta,\phi)$. Their orientation along the local surface normal is denoted by $\vec{P}$. The length of the macromolecule is $l$ and therefore the endpoint of the macromolecule simply is $\vec{r}+lR\vec{P}$, where $R$ is the rotation matrix needed to convert from the local orientation to the midpoint of the sphere (see Fig.S11a). This matrix is given by

$$R = R_z(\phi)R_y(\Theta) = \begin{pmatrix} \cos(\phi)\cos(\Theta) & -\sin(\phi) & \cos(\phi)\sin(\Theta) \\ \sin(\phi)\cos(\Theta) & \cos(\phi) & \sin(\phi)\sin(\Theta) \\ -\sin(\Theta) & 0 & \cos(\Theta) \end{pmatrix},$$

where $R_z(\phi)R_y(\Theta)$ are the usual rotation matrices along the z,y axis, respectively.
Since in the SCFS experiments the bacterium moves perpendicular to a reference surface, we use the normal of this surface to define a global z-axis. We choose the coordinates such that the origin is in the lowest point of the surface. The endpoint of the macromolecule on the sphere is $\vec{r}_1$ and the other endpoint is $\vec{r}_2$ (see Fig. S11a), such that

$$\vec{r}_1 = \vec{r} + (s+R)e_z + xe_x$$

$$\vec{r}_2 = \vec{r}_1 + lR\vec{P},$$

where $s$ is the distance from the bottom of the sphere to the reference surface (see Fig. S11). From these equations, all changes in positions, and lengths can be computed: We consider thermal length fluctuations of each macromolecule along its respective orientation $\vec{P}$. A macromolecule can bind if the closest distance to the surface is smaller than the interaction range. Once the molecule is bound it decreases its energy by potential depth $V$ which depends on the angle between the surface normal $\hat{n}$ and the molecule orientation $\vec{P}$, i.e. $V(x) = V\cos\gamma = -V\hat{n}(x)\cdot P$ (see Fig. S11b). In this attached state the molecule exerts a force onto the bacterium along $\vec{P}$, but the cantilever only measures the component along the z-direction i.e. $e_z \cdot F(l)R\vec{P}$. If the cantilever is moved the molecule is stretched and changes its orientation, such that the endpoint $\vec{r}_2$ is not moved.
Thus all coordinate changes are:

- Length fluctuations $l \mapsto l'$
    - * $\vec{r}_2 \mapsto \vec{r}_2^{'} = \vec{r}_1 + l^{'}R\vec{P}$
- Cantilever Movement: $s \mapsto s^{'}, \vec{r}_1 \mapsto \vec{r}_1^{'} = \vec{r}_1 + (s^{'}-s)e_z$
    - Molecule is not attached:
        - * $\vec{r}_2 \mapsto \vec{r}_2^{'} = \vec{r}_2 + (s^{'}-s)e_z$
    - Molecule is attached: $\vec{r}_2 \mapsto \vec{r}_2^{'} = \vec{r}_1^{'} + l^{'}RP^{'} = \vec{r}_2$
        - * $l^{'} = \sqrt{l^2 + (s-s^{'})^2 + 2l(s-s^{'})(\cos\Theta P_z - \sin\Theta P_x)}$
        - * $\vec{P}^{'} = \frac{1}{l^{'}}\begin{pmatrix} lP_x - (s-s^{'})\sin\Theta \\ lP_y \\ lP_z + (s-s^{'})\cos\Theta \end{pmatrix}$

Note that Maikranz et al. [1] use for the mechanical response of the macromolecules a WLC polymer model with probabilistic parameters. We, however, also consider situations where all molecule possess the same deterministic properties. All other considerations are identical to the model already published by Maikranz et al. [1]. Every macromolecule is placed on the surface of the sphere according to a chosen distribution and its mechanical properties can be drawn from a distribution according the publication by Maikranz et al. [1]. If not mentioned otherwise we place 50.000 molecules onto the bacterium. To increase the computation speed, macro-molecules which most likely will not come into contact with the surface are removed from the sphere. For this, the cell is placed, for every position along the substrate surface, into tangential contact of the surface (as in the geometrical model, see below), and for each macromolecules the shortest distance of the point on the sphere is computed. If this distance is larger than 70 nm, the molecule is removed.

**Computing the shortest distance to the surface** Let $(x_p, y_p, z_p)$ be the position of a point above the surface defined by $h(x,y)$. The shortest distance to the surface is then found by simple minimization of the squared distance to the surface (since the square-root is monotonic, this is equivalent but easier than minimizing the distance itself). The equations that determine the $(x,y)$ coordinates to the closest point can be derived as

$$x - x_p = (z_p - h)h_x$$

$$y - y_p = (z_p - h)h_y.$$

The shortest distance is then simply given by

$$|z_p - h(x,y)|\sqrt{h_x^2(x,y) + h_y^2(x,y) + 1}$$

Note that the equations which determine $(x,y)$ are for a sinusoidal height profile non-linear, and we there use a bisection method for computing the coordinates.

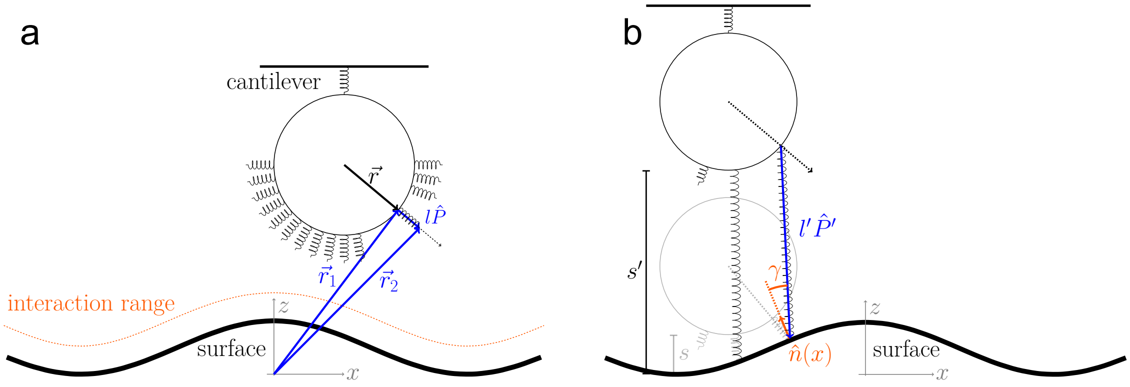


**Fig. S11 Model schematics. (a)** The macromolecules are oriented along their local surface normal given by their position $\vec{r}$ relative to the center of the bacterium. Their length $l$ and orientation $\hat{P}$ are determined by their position $\vec{r}_1$ on to the bacterium and the free moving end $\vec{r}_2$. If $\vec{r}_2$ is in the interaction range of the surface it can bind and be stretched. **(b)** If the bacterium is moved $s \mapsto s'$ perpendicular to the surface the molecules which are bound to the surface are stretched to length $l'$ and change their orientation to $\hat{P}'$. The binding strength to the surface is determined by the angle $\gamma$ between the local surface normal and the orientation $\hat{P}'$ of the macromolecule. Furthermore, the force of the molecule acts along $\hat{P}'$ such that the cantilever only measures its z-projection. Note that while the bacteria and surface dimension are scaled according to the experimental setting the spring and cantilever are not.

**Geometric Model** In this model, the relative adhesion force $f$ of the cell at point $x$ along the sinusoidal surface was computed via

$$f(x) = \frac{\sum_j w_j}{\sum_{i=1}^N w_i}.$$

Here, the j-summation runs over all rods that intersect with the surface, when the sphere was brought into tangential contact with the surface, while $w_j$ characterizes their adhesiveness. $N$ is the total number of rods on a given bacterium. Note that this model only provides relative adhesion strengths. For many rods with identical weight this model effectively computes the relative surface area of the bacterium that is able to interact with the surface. See Figure S2 for how the increasing number of rods changes the adhesion profile.

**Modified Geometric Model** In the modified geometric model we effectively incorporate angle-dependent interactions by dividing the adhesion force obtained from the geometric model by $\sqrt{1+h'(x)^2}$, where $h(x)$ parametrizes the surface. This can be justified by observing that for large distances all the macromolecules are pointing perpendicular to the surface such that $P_z \approx 1$. That this is realized in our simulations can be observed in Figure S3. Hence, the potential depth $V(x) \approx \frac{V}{\sqrt{1+h'(x)^2}}$, where we only replace the position of the individual macromolecule by the position along the surface.

#### 4 Computing the Tangential Contact Point

Every point on the bacterium's surface is parametrised by

$$\vec{B} = R \begin{pmatrix} \cos\varphi \sin\Theta \\ \sin\varphi \sin\Theta \\ \cos\Theta + 1 \end{pmatrix} + \begin{pmatrix} x \\ 0 \\ s \end{pmatrix} ,$$

where $(R, \Theta, \varphi)$ are the usual spherical coordinates, $x$ is the relative position along the surface and $s$ the distance from the minimum of the surface to the bottom of the bacterium. In order to bring the bacterium at position $x$ in tangential contact we compute the maximal $s^*$ such that

$$h(R\cos\varphi\sin\Theta + x, R\sin\varphi\sin\Theta) = R(\cos\Theta + 1) + s^* \ \forall \varphi, \Theta$$

holds. This can easily be reformulated into a maximisation problem with solution

$$s^* = \max\{h(x), h(x \pm R\sin\Theta) - R(\cos\Theta + 1)\} ,$$

where $\Theta$ needs to be determined to solve $\pm\cot\Theta\ h'(x \pm R\sin\Theta) + 1 = 0$. This $\Theta$ is the polar angle of the contact point depicted in Figure S3 and can be determined numerically.

#### 5 Substrate Parametrisation

We parametrize the surface as

$$h(x,y) = A\left(1 + \cos\left(\frac{2\pi}{\lambda}x\right)\right)$$

with $A = 190$ nm , $\lambda = 2750$ nm matching the experimentally used dimensions. Note that the bacterium has a radius of 500 nm.

#### 6 Supplementary Table

**Table S1 Experimental parameters for all cells.** The table gives for all cells in Figure 3 and 4 of the main text the total number of consecutive force-distance curves, the x-offset between two neighboring curves and the used sample.

| cell no. | 1 | 2 | 3 | 4 | 5 | 6 | 7 | 8 | 9 | 10 | 11 | 12 | 13 | 14 |
|---|---|---|---|---|---|---|---|---|---|---|---|---|---|---|
| # of curves | 500 | 500 | 500 | 400 | 400 | 400 | 500 | 450 | 500 | 450 | 400 | 400 | 500 | 500 |
| x-offset [nm] | 20 | 20 | 20 | 25 | 25 | 25 | 20 | 30 | 20 | 30 | 25 | 25 | 20 | 20 |
| sample no. | 2 | 2 | 2 | 1 | 1 | 1 | 2 | 3 | 2 | 3 | 1 | 1 | 2 | 2 |

#### Notes and references

1 E. Maikranz, C. Spengler, N. Thewes, A. Thewes, F. Nolle, P. Jung, M. Bischoff, L. Santen and K. Jacobs, *Nanoscale*, 2020, **12**, 19267–19275.
